# The fungal RNA-binding protein Ssd1 represses Sun4 protein abundance through recognition of 5ʹ UTR structural and sequence elements

**DOI:** 10.64898/2026.08.03.741543

**Authors:** Benjamin Kleinerman, Uma Jayachandran, Marah M. Jnied, Weronika A. Danecka, Aleksandra Rosinska, Edward W.J. Wallace, Atlanta G. Cook

## Abstract

Regulation of protein abundance allows fungi to adapt to changing environments, regulate their growth and morphology, and react to external stresses. Ssd1 is a fungal RNA binding protein that binds mRNAs encoding cell wall remodelling proteins and regulates cell wall biogenesis. Ssd1 binding sites (SBSs) have been described by computational and biochemical analyses, but how Ssd1 recognises these sites and how that relates to regulation of protein abundance was less clear. Here, a co-crystal structure of *Saccharomyces cerevisiae* Ssd1 with an SBS reveals core determinants of recognition, while fluorescent reporters of Sun4, an Ssd1-regulated cell wall protein, were used to characterise structure- guided mutations. We find that Ssd1 has an extensive RNA binding site that recognises two elements of the SBS: an upstream element that forms a structural motif, and an element containing tandem CNYU sequences that engages Ssd1 in base-specific recognition. Mutations to the Ssd1 RNA binding surface prevent Ssd1-dependent repression of fluorescent reporters and show strong phenotypes in assays of cell wall stress resistance and genetic interactions with the Cbk1 kinase. Loss of repression is also observed if both *SUN4* SBSs are altered. However, the presence of one SBS in the *SUN4* 5ʹ untranslated region is sufficient to confer Ssd1-dependent suppression of protein abundance. Our work confirms that RNA binding is a core function of Ssd1 and is likely to inform functional analyses in fungi beyond *S. cerevisiae*, where Ssd1 orthologs have been identified as virulence factors in several fungal pathogens.

## Introduction

Regulation of protein abundance is critical to the survival of microorganisms, including fungi, allowing them to react appropriately to environmental stimuli and stresses. *Saccharomyces cerevisiae* (*Sc*) Ssd1 is an RNA binding protein, conserved in fungi, that regulates expression of cell wall proteins through post-transcriptional mechanisms (Ballou *et al*, 2021; Bayne *et al*, 2021; Bresson *et al*, 2020; Hall & Wallace, 2022; Jansen *et al*, 2009; Kaeberlein & Guarente, 2002; Uesono *et al*, 1997; Wanless *et al*, 2014). In response to a variety of stress conditions, *Sc*Ssd1 shows substantial changes in its RNA association (Bresson *et al*., 2020). While Ssd1 is not essential in *S. cerevisiae*, its loss can have pleiotropic effects and, consequently, it has been identified in several genetic screens (Luukkonen & Seraphin, 1999; Stettler *et al*, 1993; Sutton *et al*, 1991; Tung *et al*, 2021; Uesono *et al*, 1994; Wilson *et al*, 1991; Xu *et al*, 2020). In wild-living *S. cerevisiae* strains, Ssd1 is needed for tolerance of naturally occurring aneuploidy, through a poorly understood mechanism (Dutcher & Gasch, 2024; Dutcher *et al*, 2024; Hose *et al*, 2020). In other fungi, Ssd1 contributes to processes associated with cell wall remodelling, such as cell separation in *Cryptococcus neoformans* Titan-inducing growth conditions and hyphal transport of ribonucleoprotein particles in *Aspergillus nidulans* (Ballou *et al*., 2021; Modaffari *et al*, 2026). As dynamic cell wall remodelling is important in pathogenic fungi, Ssd1 has been identified as a virulence factor in both animal and plant fungal pathogens (Gank *et al*, 2008; Hall & Wallace, 2022; Schwarzmuller *et al*, 2014; Thammahong *et al*, 2019). A deeper understanding of Ssd1 association with RNA would allow dissection of its role in fungal virulence.

Prior analyses to identify mRNAs associated with Ssd1 found that high confidence mRNA targets encode cell wall remodelling proteins, such as members of the SUN family of proteins, including Sun4, Uth1 and Sim1 (Bayne *et al*., 2021; Hogan *et al*, 2008). Sun4 plays important roles in bud scar formation and septum remodelling and is typically localized to the endoplasmic reticulum, the cell periphery, the cell septum and bud scars (Ast *et al*, 2013; Firon *et al*, 2007; Kuznetsov *et al*, 2016; Mouassite *et al*, 2000). Prior work has also identified transcripts of the cell cycle regulator *CLN2* as a target of Ssd1 (Hogan *et al*., 2008; Ohyama *et al*, 2010). Ssd1 acts downstream of Cbk1 kinase, a master regulator of cell growth and morphogenesis. When Cbk1 activity is lost or blocked, the presence of wild type Ssd1 causes growth arrest and cell death (Du & Novick, 2002; Jansen *et al*., 2009).

Based on the identification of an Ssd1-dependent motif enriched in untranslated regions (UTRs) of mRNA targets (Hogan *et al*., 2008), prior studies characterized the role of 5ʹ and 3ʹ UTRs in Ssd1-dependent expression of targeted proteins and showed that Ssd1 could immunoprecipitate UTR sequences, suggesting direct binding (Ohyama *et al*., 2010; Wanless *et al*., 2014). It was also shown that Ssd1 could suppress translation of associated transcripts (Jansen *et al*., 2009; Ohyama *et al*., 2010; Wanless *et al*., 2014). However, a coherent model of how RNA binding activity of Ssd1 determined protein abundance was hampered by an incomplete description of the Ssd1 target sequence (Hogan *et al*., 2008; Wanless *et al*., 2014).

In previous work, we identified *S. cerevisiae* Ssd1 binding sites (SBSs) at nucleotide resolution using UV crosslinking and analysis of cDNAs and found the strongest enrichments in 5ʹ UTRs (Bayne *et al*., 2021). Computational analysis of recurring motifs in bound mRNAs revealed two elements that contribute to binding. An upstream element (UE) occurs in a subset of highly bound sites. A conserved downstream element with the consensus sequence CNYUCNYU (N is any base, Y is a pyrimidine) is similar to the previous description of a common element (Hogan *et al*., 2008). We refer to the downstream element as the CNYU element (CE). CEs are identifiable in transcripts encoding cell wall proteins in many fungal species (Bayne *et al*., 2021; Modaffari *et al*., 2026; Schrettenbrunner *et al*, 2026). For *Sc*Ssd1, neither the UE nor the CE individually bind well to Ssd1 and only SBSs that encompass both elements bind with high affinity *in vitro* (Bayne *et al*., 2021).

Here we focus on the regulation of Sun4 by Ssd1 in *S. cerevisiae*. Using *in vivo* reporter assays, we show that regulated repression of Sun4 depends on *cis*-acting sequences in the 5ʹ UTR of the *SUN4* transcript. We report a co-crystal structure of Ssd1 with a *SUN4* SBS, which reveals a novel mode of protein-RNA binding, in which Ssd1 directly recognises both elements of a cognate SBS, using structure- and sequence-specific recognition. We further show that repression of Sun4 protein abundance depends directly on RNA recognition by Ssd1, and that a single SBS at the native *SUN4* locus can mediate repression.

## Results

### Ssd1 binding sites in the SUN4 5ʹ UTR are needed to repress Sun4 protein abundance

To gain insight into how Ssd1 regulates a specific mRNA, we focused on the *SUN4* gene that we previously identified as a high confidence cellular target of Ssd1 (Bayne *et al*., 2021). Two SBS sequences are strongly enriched for Ssd1 in the 5ʹ UTR of *SUN4*, with smaller peaks across the coding sequence (CDS) and 3ʹ UTR, including at two CDS-located SBSs (Fig. 1a). Multiple SBSs are also found in 5ʹ UTRs of *SUN4* homologs from other ascomycete fungi (Bayne *et al*., 2021). *SUN4* has an intron in its 5ʹ UTR, which is rare amongst *S. cerevisiae* genes, but no SBSs are found in the intron sequence (Fig. 1a).

**Figure 1.**
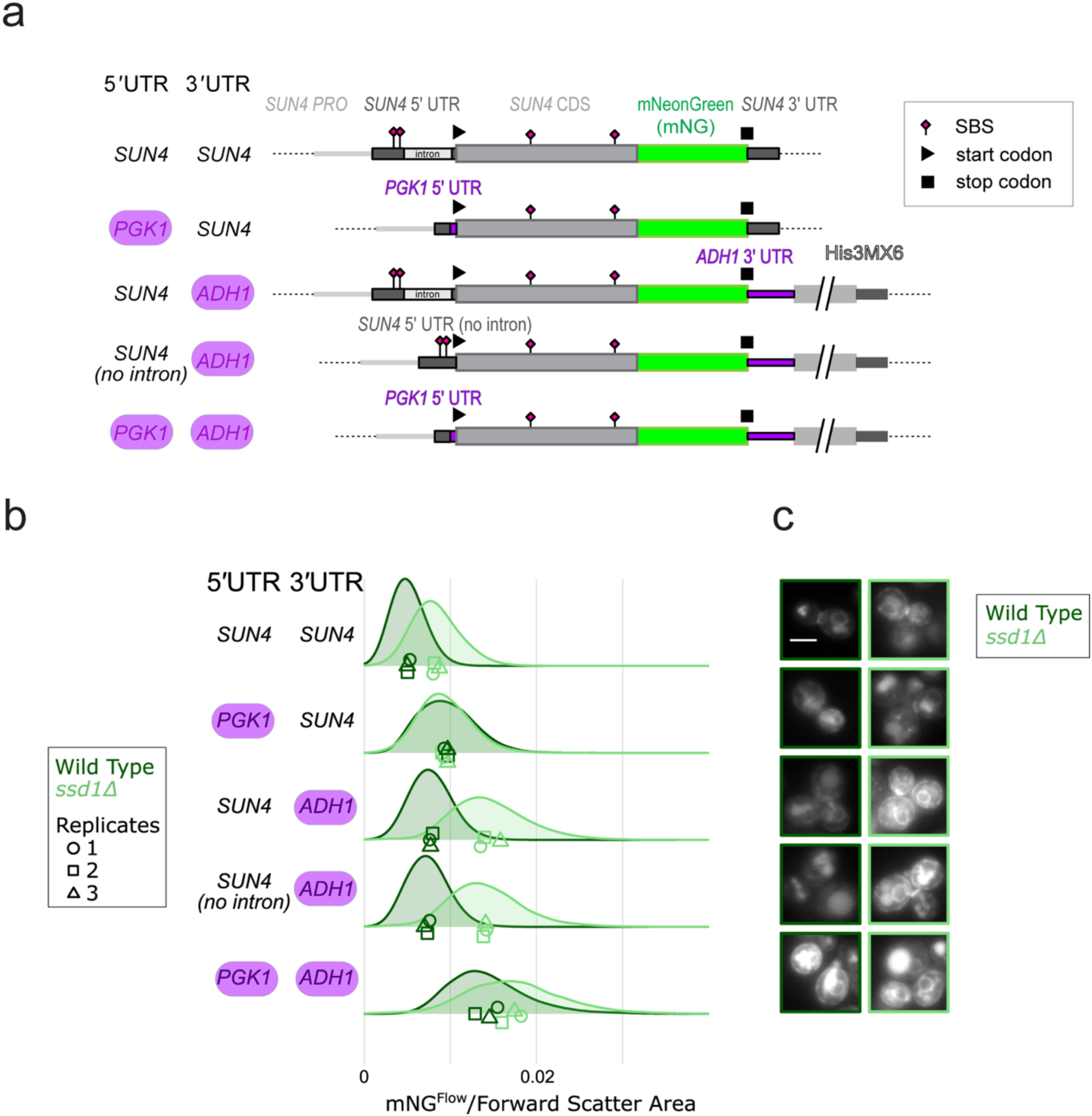
The 5ʹ UTR of *SUN4* is needed to repress Sun4 protein expression. (a) Schematic overview of *SUN4* constructs tagged with mNeonGreen (*SUN4-mNG*) at the native yeast *SUN4* locus, some with UTRs replaced by alternative yeast UTRs (from *PGK1* and *ADH1*) lacking Ssd1 binding sites (SBSs). Native *SUN4* parts are in grey and green, exogenous parts are in purple, and a yeast selectable marker cassette (HIS3MX6) is in grey. Thicker boxes indicate coding sequences (CDS), and thinner boxes indicate untranslated regions (UTRs), with internal lighter grey indicating a native 5ʹ UTR intron. Pink diamonds indicate SBSs; start and stop codons are indicated. (b) Abundance of Sun4-mNG and variants with replaced UTRs, in WT and *ssd1Δ* backgrounds, measured by flow cytometry. All flow cytometry data are reported in arbitrary units; mNG signal is divided by forward scatter area to account for cell size. Three biological replicates, from independent single colonies, were measured and plotted as a single distribution, and their means are plotted individually. (c) Fluorescence microscopy images of Sun4-mNG and variants, showing green fluorescence signal. The scale bar in the upper left image indicates 5 µm. To ensure comparability, all images shown were taken on the same day with identical microscope settings and processed identically.

To characterise Ssd1 regulation of Sun4 protein abundance, we fused the coding sequence of mNeonGreen (mNG) in-frame with the Sun4 coding sequence at its native locus, to encode a C-terminally tagged protein. We then quantified green fluorescence of yeast cells by flow cytometry, in a wild-type or *ssd1*Δ background, correcting for cell size after gating for singlet cells (Fig. S1a,b). As expected, when *SUN4-mNG* retains its native *SUN4* UTRs, Sun4-mNG protein shows Ssd1-dependent abundance: green fluorescence is lower in the wild-type background compared to *ssd1Δ* cells (Fig. 1b). To dissect which UTR sequences contribute to Ssd1-dependent regulation, we generated a series of strains where we replaced the native *SUN4* 5ʹ, 3ʹ, or both UTRs with other *S. cerevisiae* UTRs that have no SBSs (Fig. 1a). Replacing the *SUN4* 5ʹ UTR with the *PGK1* 5ʹ UTR removes the dependence on *SSD1* (Fig. 1b). In contrast, replacing the *SUN4* 3ʹ UTR with the *ADH1* 3ʹ UTR retains Ssd1 regulation (Fig. 1b). Removing the 5ʹ UTR intron in this context does not change Ssd1 dependency nor the overall amount of fluorescent protein expressed. Lastly, replacing both UTRs shows loss of Ssd1 dependence of Sun4-mNG abundance in cells, showing that the 5ʹ UTR is the primary site of regulation. Of note, all strains that have non-native 3ʹ UTRs have higher fluorescence regardless of whether Ssd1-dependent regulation is present (Fig. 1b). This is consistent with previous reporter gene assays showing that the *SUN4* 3ʹ UTR leads to low protein production relative to a set of 10 yeast 3ʹ UTRs (Haynes *et al*, 2022).

Fluorescence levels observed by flow cytometry are consistent with fluorescence microscopy (Fig. 1c). Sun4-mNG localization has a similar pattern to prior analyses that showed endoplasmic reticulum and cell periphery localization, consistent with its known functions (Ast *et al*., 2013; Dubreuil *et al*, 2019; Firon *et al*., 2007; Kuznetsov *et al*., 2016; Mouassite *et al*., 2000). Although deleting *SSD1* or replacing the UTRs of *SUN4* increases the abundance of Sun4-mNG, we see no obvious differences in its localization.

A possible explanation for the Ssd1-mediated repression of Sun4-mNG abundance is that Ssd1 reduces *SUN4* mRNA abundance. RT-qPCR performed on the same set of native and variant *SUN4-mNG* constructs showed variable levels of *SUN4-mNG* mRNAs across all experiments that did not match the changes observed by flow cytometry (Fig. S1c). When fluorescence is compared to *SUN4-mNG* mRNA abundance for all UTR variants, protein abundance was uncorrelated with differences in mRNA abundance (Fig. S1d). This indicates that the effects of Ssd1 on Sun4-mNG protein abundance are not driven by changes in *SUN4-mNG* mRNA abundance.

Our data show that Ssd1 mediates repression of Sun4 protein abundance primarily through *cis*-acting sequences in the *SUN4* 5ʹ UTR. In contrast, other sequences in the transcript (3ʹ UTR, 5ʹ intron and the CDS) contribute little to Ssd1-mediated repression of Sun4. We therefore focused on how Ssd1 recognises SBS sequences of the *SUN4* 5ʹ UTR.

### The crystal structure of Ssd1 with an RNA binding motif from *SUN4* 5ʹ UTR

To understand how Ssd1 recognises SBSs, we co-purified (Fig. S2a) and co-crystallised Ssd1 with a 26-nucleotide RNA derived from the *SUN4* 5ʹ UTR, here called *SUN4duo*, that encompasses both a UE and a CE. As the N-terminal region of Ssd1 is natively unstructured, we used a previously characterized Ssd1ΔN338 construct for co-crystallisation (Fig. 2a) (Bayne *et al*., 2021). This fragment includes two identifiable cold shock domains (CSD1 and CSD2), an RNAse II-like domain (RNB) and an S1 domain. These four domains together form a single structural unit. Crystals of Ssd1ΔN338-*SUN4duo* diffracted to 2.5 Å, in space group *P*2_1_2_1_2_1_ (Table 1). After molecular replacement with coordinates of Ssd1ΔN338 (PDBid 7AM1), clear difference density was observed for *SUN4duo* over an extended surface of the two CSD domains of Ssd1. The RNA structure was built into the map and completed through iterative model building and refinement. The final model has R_work_ and R_free_ values of 21 % and 27% respectively, with good stereochemistry (Table 1).

**Figure 2.**
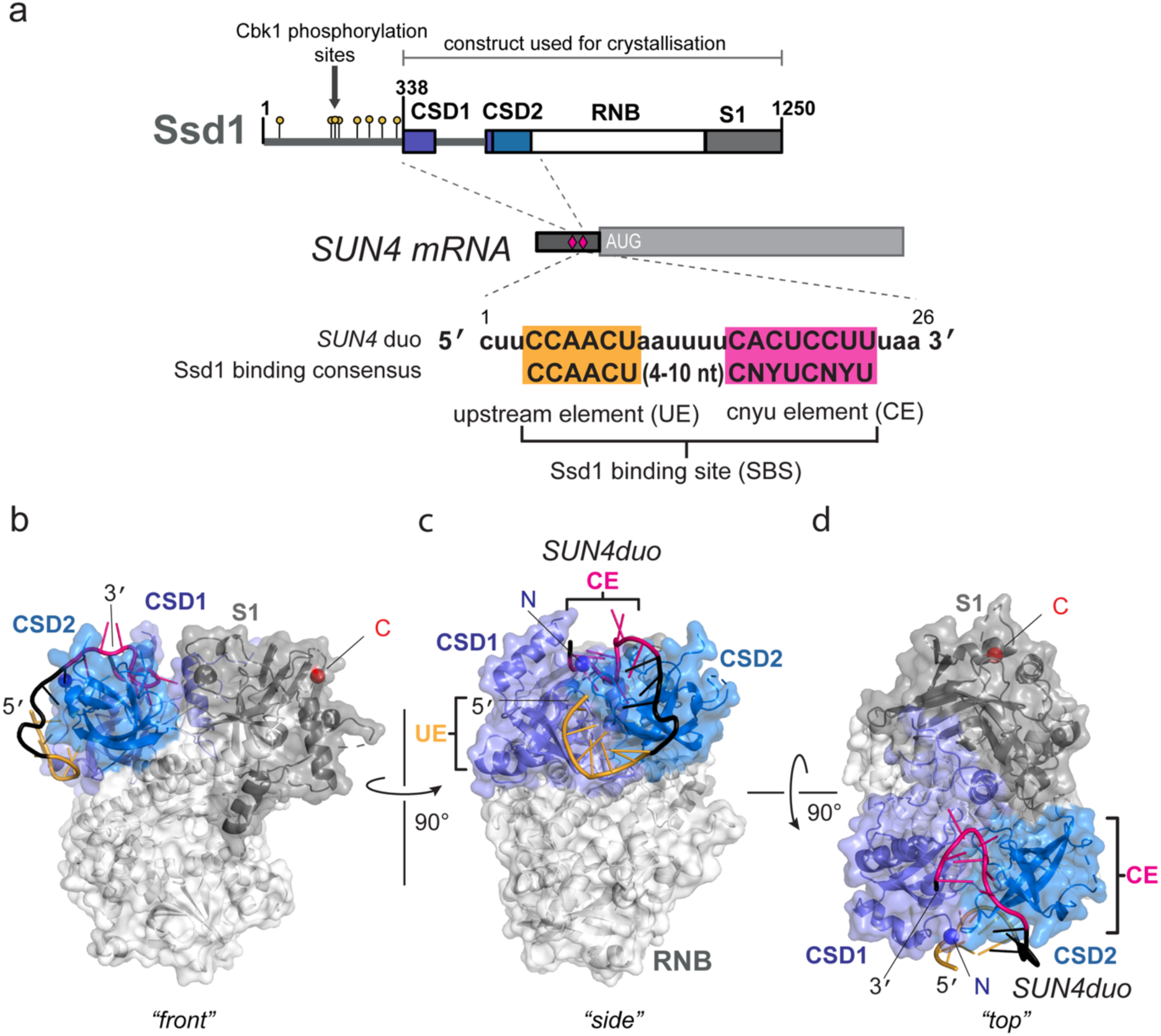
Structural overview of Ssd1-*SUN4duo* complex. (**a**) Schematic of Ssd1 and *SUN4* 5ʹ UTR. The start codon is represented by “AUG”. Pink diamonds indicate SBSs. An expanded view of the SBS used in crystallization is shown at the bottom, with core elements highlighted in orange and hot pink. (**b**) Cartoon and surface rendering of the “front” view of the Ssd1-*SUN4duo* complex, with CSD domains on the top left and the C-terminal S1 domain at the top right. The RNA is shown as cartoon with UE in orange, linker in black, and CE in hot pink. (**c**) “Side” view of the UE motif bound to CSD domains of Ssd1, using a vertical rotation of 90° with respect to (b). (**d**) View of the CE binding to Ssd1 along the “top” surface of Ssd1, using a rotation around the horizontal axis compared to (c). Blue and red spheres indicate the N and C terminus of Ssd1, respectively.

**Table 1.**
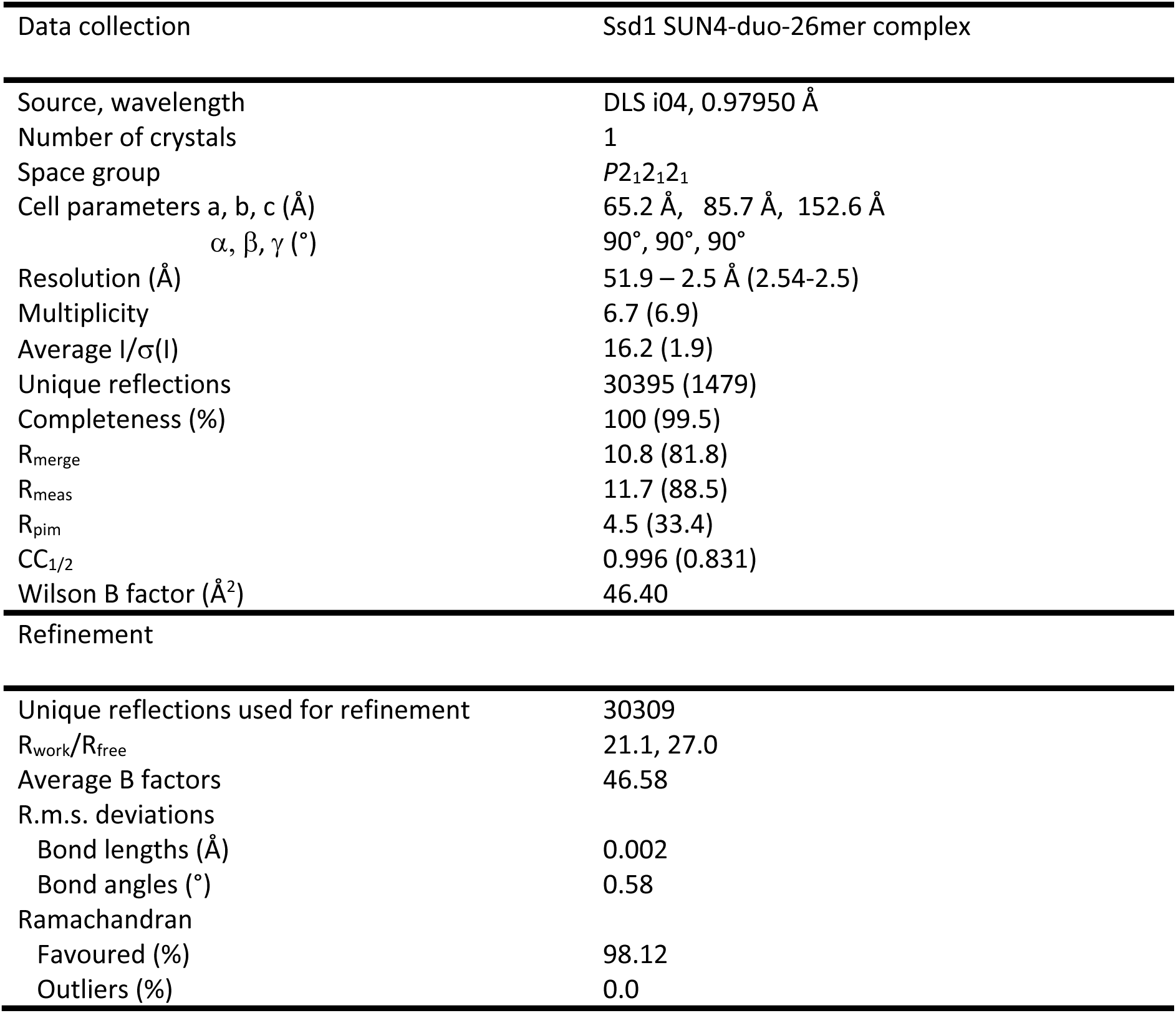
xxxxx

| Data collection | Ssd1 SUN4-duo-26mer complex |
| --- | --- |
| Source, wavelength | DLS i04, 0.97950 Å |
| Number of crystals | 1 |
| Space group | *P*2<sub>1</sub>2<sub>1</sub>2<sub>1</sub> |
| Cell parameters a, b, c (Å) | 65.2 Å, 85.7 Å, 152.6 Å |
| $\alpha, \beta, \gamma$ (°) | 90°, 90°, 90° |
| Resolution (Å) | 51.9 – 2.5 Å (2.54-2.5) |
| Multiplicity | 6.7 (6.9) |
| Average I/$\sigma$(I) | 16.2 (1.9) |
| Unique reflections | 30395 (1479) |
| Completeness (%) | 100 (99.5) |
| R<sub>merge</sub> | 10.8 (81.8) |
| R<sub>meas</sub> | 11.7 (88.5) |
| R<sub>pim</sub> | 4.5 (33.4) |
| CC<sub>1/2</sub> | 0.996 (0.831) |
| Wilson B factor (Å<sup>2</sup>) | 46.40 |
| Refinement |  |
| Unique reflections used for refinement | 30309 |
| R<sub>work</sub>/R<sub>free</sub> | 21.1, 27.0 |
| Average B factors | 46.58 |
| R.m.s. deviations |  |
| Bond lengths (Å) | 0.002 |
| Bond angles (°) | 0.58 |
| Ramachandran |  |
| Favoured (%) | 98.12 |
| Outliers (%) | 0.0 |

When compared with Ssd1 in the absence of RNA (Bayne *et al*., 2021), the overall structure of the protein is very similar, with a root mean square displacement (r.m.s.d) on all Cɑ = 0.79 Å (Fig. S2b). Superposition of the CSD domains (r.m.s.d. = 0.74 Å over Cɑ residues 341-642) reveals small changes to loop structures (Fig. S2c). However, for the RNB and S1 domains together, the r.m.s.d. is 0.52 Å (over Cɑ residues 642-1248), suggesting that there are no substantive structural changes to these domains. Rather, the differences can be explained by a rigid rotation of the CSD domains towards the core of the protein (Fig. S2d). Two segments of Ssd1 that were not visible in electron density maps of the apo Ssd1 structure could be built in the presence of RNA, including the backbone residues of an ɑ-helix extending from residues 415-423 and a segment of an extended loop encompassing residues 562-566. This suggests that RNA induces some local disorder-to-order transitions in Ssd1.

### A surface extending over the Ssd1 cold shock domains binds to *SUN4duo*

The *SUN4duo* RNA binding site extends over the two CSDs, burying 1892 Å^2^ of the Ssd1 surface. We previously showed that this surface is highly conserved and has an overall basic charge (Ballou *et al*., 2021; Bayne *et al*., 2021). Nucleotides 4-23 of the RNA could be built (Fig. 2b, Fig. S2e,f), although only the backbone was visible for A11 and U12. The 5ʹ UE, encompassing the CCAACUA sequence (nucleotides 4-10 inclusive), binds at the interface between the CSDs, close to where these two domains meet the RNB domain (Fig. 2c). These seven nucleotides describe a curved loop, whereby C4-A6 interact with CSD1, then the backbone of the RNA changes direction between A6 and A7 such that A7-A10 interact with the Ssd1 at the junction between CSD1 and CSD2. A11-U16 are part of a variable linker sequence between the UE and CE sequences that extends over the surface of CSD2, towards the “top” of Ssd1 where the CE binds (Fig. 2c,d). The backbone of the nucleotides that make up the CE extend along the top surface of Ssd1 with a tight turn sending the backbone back towards the beginning of the CE, splaying bases into pockets on the Ssd1 surface (Fig. 2d). The extensive interactions of the SBS with Ssd1 are consistent with the high affinity binding (Kd = 8 nM) of the RNA (Bayne *et al*., 2021).

### The SUN4 upstream element is a structural motif

The UE forms an unusual structural motif with stacking interactions between bases C4-A6. A Mg^2+^ ion mediates the abrupt turn in the backbone between the phosphate groups of A6 and A7 that creates an antiparallel base stack formed by A7-U9 (Fig. 3a,b, Fig. S3a). A10 caps off this structure by stacking across neighbouring A6 and U9 bases (Fig. 3b). No Watson-Crick base pairs form between these two stacks. Instead, a hydrogen bonding network connects different base moieties to stabilise the structure (Fig. 3b, Fig. S3a,b). For example, the Hoogsteen edge of A7 forms two hydrogen bonds with the o2 and o2ʹ atoms of C5 (Fig. S3b). The o2 atom of C5 also hydrogen bonds to n4 of C8, while the n4 amine moiety of C5 is within hydrogen bonding distance of the n6 amine of A6 (Fig. 3b). The U9 o4 carbonyl is in proximity to the n6 amine of A10 that also connects with the ribose o2ʹ of A6 (Fig. S3a). These interactions connect the bases in a single structural motif.

**Figure 3.**
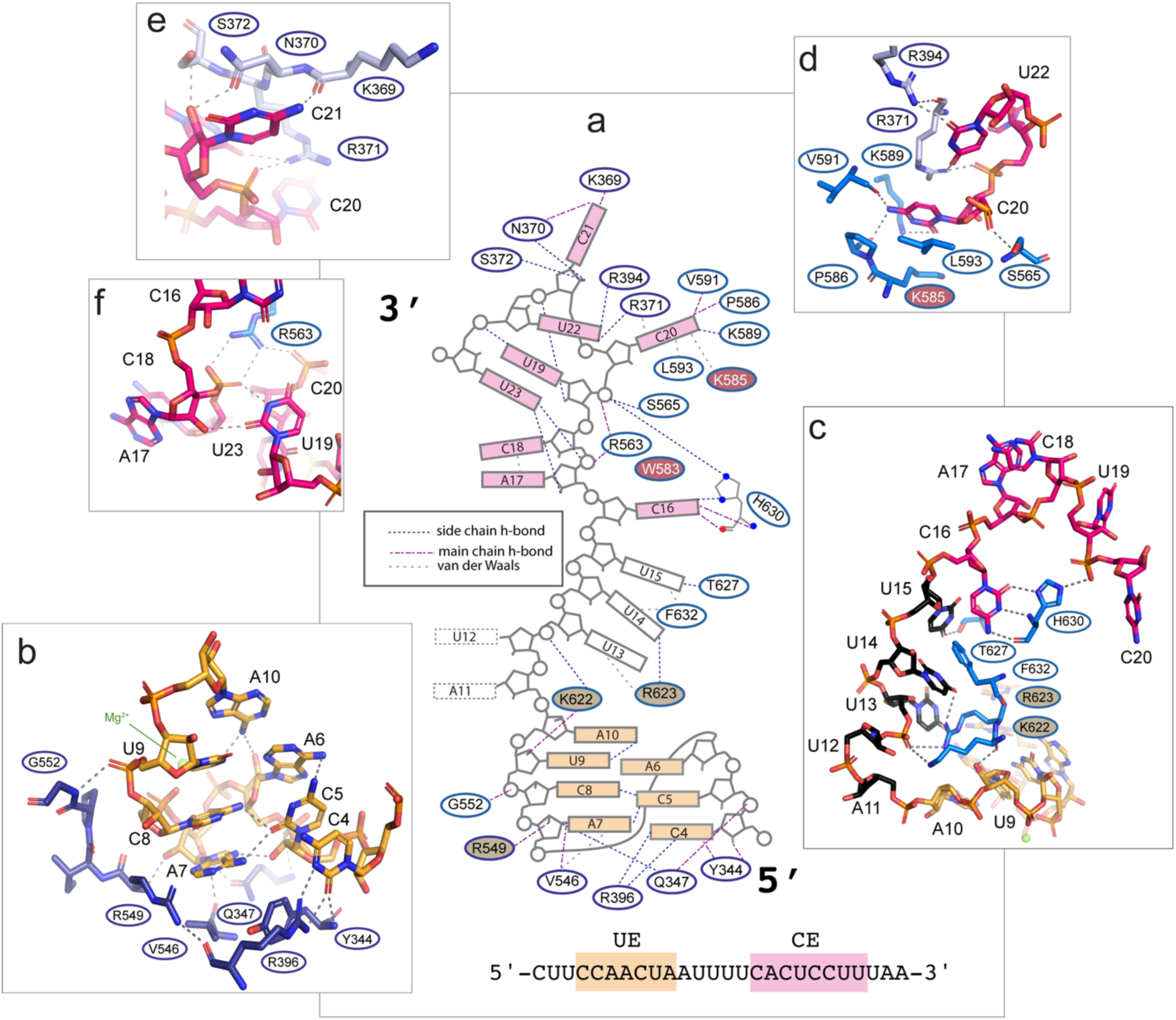
The SBS is recognized through both structure- and sequence-specific interactions. (**a**) Schematic overview of *SUN4duo* SBS motif binding to Ssd1. UE bases are colored pale orange, and CE bases are colored pink. Ovals indicate Ssd1 residues that interact with RNA. Residues with filled ovals indicate sites that were previously shown to affect RNA binding when mutated. The key indicates different types of interactions denoted by dotted lines. (**b**) Interactions between UE and Ssd1. For panels b-f, structures are shown as sticks and gray dotted lines indicate hydrogen bonding interactions. (**c**) Interactions between the linker region nucleotides and Ssd1. (**d**) The C20 and U22 binding pocket. (**e**) C21 interactions with Ssd1. (**f**) The latch closure of U23.

As the bases of the UE mainly interact with each other, there are few base-specific interactions with Ssd1. One direct recognition event occurs with C4, which makes a bipartite hydrogen bond via its Watson-Crick (WC) edge (n3 and o2), with Arg396 (Fig. 3b, S3c). The position of C4 is further stabilised by a mainchain interaction of Tyr344 to o2. This residue also forms a π- π interaction with Arg396 (Fig. S3c). Mainchain and side chain interactions of Gln347, Val546, Arg549 and Gly552 with ribose and phosphate moieties further stabilise the UE sequence on the surface of Ssd1 (Fig. 3b).

### Base-specific interactions define the CNYU element

Prior bioinformatic analyses found variable lengths (4-10 nt) of the linker region that connects UE to the CE (Bayne *et al*., 2021). Bases 11 and 12 are not well defined in the electron density map and only backbone moieties could be built (Fig 3a,c), suggesting that additional bases could be accommodated at this point. In contrast, U13-U15 are well defined in the electron density and interact with K622, R623 and F632 as the linker traverses the CSD-side surface to the CSD-top surface (Fig. 3c). Mutations to K622 and R623 were previously shown to disrupt SBS RNA binding (Bayne *et al*., 2021). Two direct base interactions are observed between U14 and R623 and between U15 and the hydroxyl side chain of T627 (Fig. 3c).

The first base of the CNYU element, C16, is exquisitely recognised through its WC edge by hydrogen bonds across the sidechain and mainchain of H630 (Fig. 3c). This explains the strong preference for a C base at the first position of the **C**NYUCNYU consensus motif. H630 also connects to the backbone phosphate moiety of C20, bridging the beginning and middle of the CE. A17 and C18 (C**NY**UCNYU) are flipped away from the surface of the protein and stack against each other, explaining why most bases can be accommodated at these positions (Fig. 3c, Fig S3d). The flipped out conformation is stabilised by R563 and W583 (Fig S3d), the latter of which was previously identified as contributing to RNA binding (Bayne *et al*., 2021). U19, at position 4 (CNY**U**CNYU) plays a structural role in positioning the RNA by connecting to the backbone moieties of U22 (via o2ʹ) and U23 (n3 bridging via a crystallographic water molecule to U23 phosphate) (Fig. S3d). There are no base-specific interactions observed for U19; the strong preference for uridine bases at this position may relate to the smaller size of pyrimidine bases and the less polar nature of uridine compared to cytosine.

C20, at position 5 (CNYU**C**NYU), is bound in a deep pocket on the Ssd1 surface made up of residues K585, P586, K589, V591 and L593 (Fig. 3d). Base-specific interactions include two n4 hydrogen bonds with the carbonyl groups of P586 and V591 and an o2 hydrogen bond with the side chain of K589. These interactions probe specific donor and acceptor groups of the WC edge of this base, explaining the strong requirement for C at this position. A π-π interaction with R371 connects this binding pocket to a neighbouring binding site for U22 (CNYUCN**Y**U) (Fig. 3d). The o2 and o4 carbonyl moieties of U22 accept hydrogen bonds from R371 and R394, suggesting a strong selection for U at this position. Indeed, although this position is described as preferential for pyrimidines, uridines are most frequently found at this site (Bayne *et al*., 2021). The intervening nucleotide, C21 (CNYUC**N**YU), bound by K369, N370, and S372, is more surface exposed than C20 or U22, consistent with non-specificity for this position (Fig. 3e). The final nucleotide of the element, U23 (CNYUCNY**U**), acts as a structural latch that closes the splayed structure of the CE by interacting with the A17 ribose and C18 phosphate backbone moieties (Fig 3f). The U23 interactions use the WC base-specific pattern of hydrogen bond donor (n3) and acceptor (o2) moieties. This structural connection is reinforced by R563 interactions with the C18 and C20 phosphate groups. In summary, a combination of intramolecular RNA structural interactions and base-specific interactions with Ssd1 residues explains the recognition of the CNYUCNYU element.

### Ssd1 RNA recognition point mutants show loss-of-function phenotypes

To test the consequences of Ssd1-RNA recognition for cell growth and RNA regulation, we made a series of mutations to the RNA binding surface of Ssd1 (Fig. 4a) and tested their effects in three assays. First, we tested the ability of Ssd1 and point mutants to promote resistance to cell wall stress induced by the chitin binding molecule calcofluor white (CFW). We mutated *Sc*Ssd1 C-terminally tagged with a His6-FLAG tag (Ssd1-HF) at its native locus, as described previously (Bayne *et al*., 2021). Wild-type Ssd1 or Ssd1-HF support growth on plates containing 25 µM CFW, but *ssd1Δ* does not (Fig. 4b). Ssd1 point mutants all have reduced resistance to cell wall stress compared with wild-type cells, with the most severe phenotypes seen for R394A, W583A, and H630A. All strains grow well in the absence of cell wall stress, and Ssd1 mutant proteins are similarly abundant to WT Ssd1-HF as shown by western blotting (Fig. S4a). This shows that Ssd1 protein function, not abundance, accounts for the cell wall stress phenotypes observed in point mutants.

**Figure 4.**
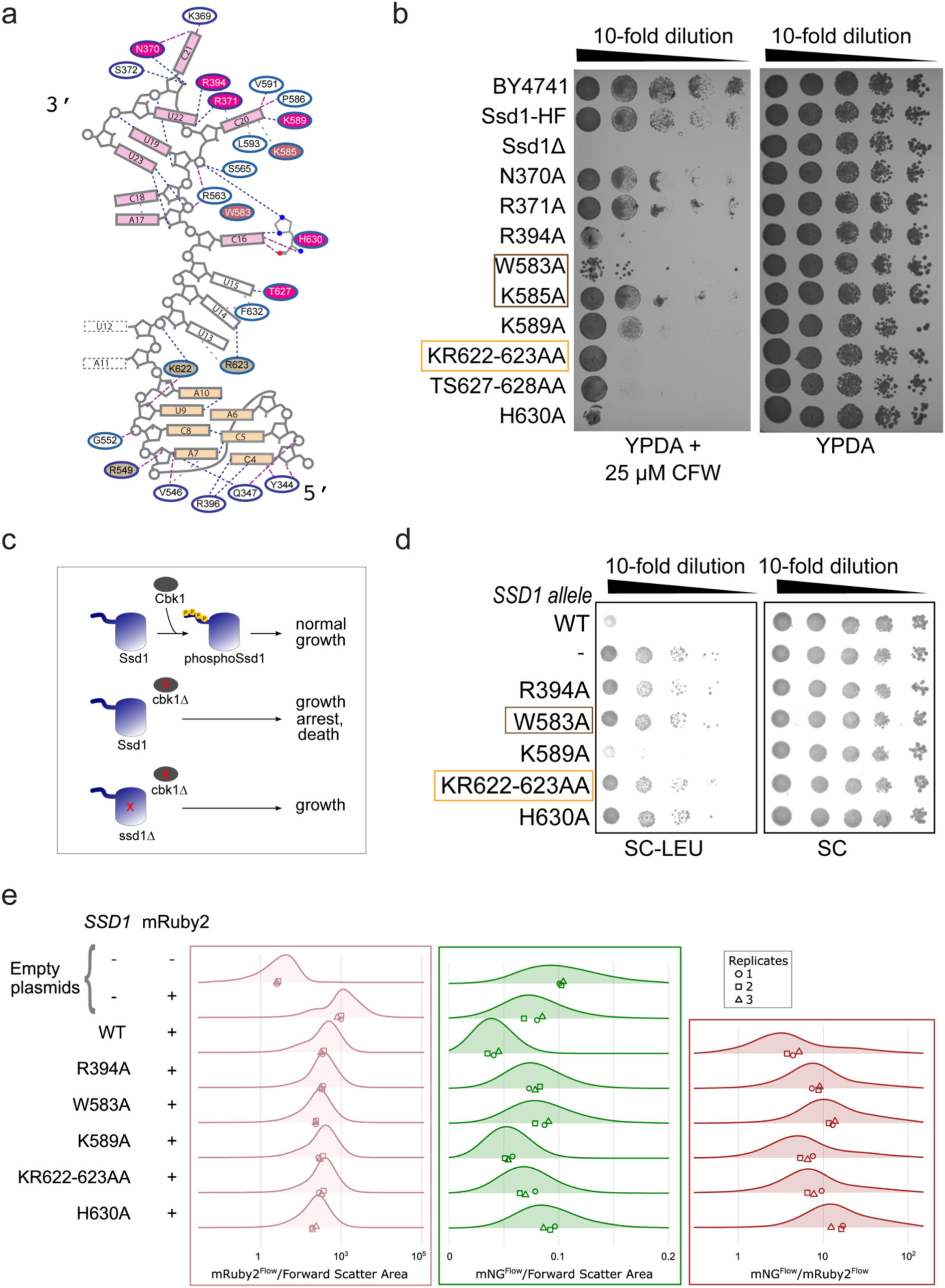
Point mutations to the Ssd1 RNA-binding surface cause loss of function phenotypes. (**a**) Overview of Ssd1 point mutations based on the schematic in Fig. 3a. (**b**) Cell wall stress sensitivity assayed by 10-fold serial dilutions of yeast strains spotted and grown at 30°C on YPDA with (left) or without (right) 25 µM calcofluor white (CFW). Wild type is BY4741; Ssd1-HF has a C-terminal His-FLAG tag; *ssd1Δ* is a deletion mutant; individual and paired Ssd1-HF RNA-binding point mutants are indicated. Boxed are residues previously characterized as “Ssd1-top” (brown) and “Ssd1-side” (orange) mutations. (**c**) Schematic of functional interactions between Cbk1 and Ssd1. Phosphorylation of Ssd1 by Cbk1 is needed for normal growth. Loss of Cbk1 prevents cell wall growth, leading to arrest and eventually death. Loss of both Cbk1 and Ssd1 supports growth. (**d**) LEU-selectable plasmids encoding WT *SSD1,* or variants, fused to mRuby2 and expressed from the yeast *RNR2* promoter, were transformed into a *cbk1Δ ssd1Δ* background. 10-fold serial dilutions of transformation mixtures were spotted and grown at 30°C on yeast synthetic media without or with leucine. Empty plasmid control (-) expresses only mRuby2 (no *SSD1*). (**e**) Abundance of plasmid- expressed Ssd1-mRuby2 RNA-binding point mutants (pink), Sun4-mNG (green), and the ratio of Sun4-mNG to Ssd1-mRuby2 (brown), measured by flow cytometry. Ssd1-mRuby2 fluorescence is displayed on a log scale, Sun4-mNG fluorescence on a linear scale, and their ratio on a log scale. Two empty plasmid controls are included: no coding sequence expressed and a plasmid expressing only mRuby2. Three biological replicates, from independent single colonies, were measured, and their means are plotted.

Second, focusing on a smaller subset of residues involved in base recognition, we tested the ability of Ssd1 and point mutants to compromise *S. cerevisiae* growth in the absence of the kinase Cbk1. It was previously observed that *cbk1Δ* cells expressing full-length Ssd1 are inviable, but *cbk1Δ ssd1Δ* cells are viable (Du & Novick, 2002). Because inhibition of Cbk1 blocks bud growth (Kurischko *et al*, 2008), it is likely that the lack of Cbk1-mediated phosphorylation of Ssd1 arrests growth (Fig. 4c). We hypothesised that mutations that prevent RNA recognition by Ssd1 would relieve this growth arrest. To test this, we complemented a *cbk1Δ ssd1Δ* strain with centromeric plasmids encoding an auxotrophic selection marker for growth on leucine-containing media (SC-LEU) and an expression cassette for *SSD1* or mutant alleles, fused in-frame to the mRuby2 fluorescent marker. As expected, plasmids expressing wild-type Ssd1 prevent growth in SC-LEU while a plasmid lacking any *SSD1* allele confers growth in SC-LEU (Fig. 4d). Plasmids expressing Ssd1 point mutants R394A, W583A, KR622-623AA, and H630A confer growth in SC-LEU, indicating that the Ssd1 function relevant to genetic interactions with Cbk1 is lost. By contrast, a plasmid expressing Ssd1-K589A does not confer growth in SC-LEU, indicating a weaker disruption of the wild-type function.

Third, we tested the ability of Ssd1 and point mutants to repress Sun4-mNG abundance in our flow cytometry assay. We transformed the above plasmids, expressing Ssd1-mRuby2 variants, into *CBK1 ssd1Δ* cells that express Sun4-mNG with a *SUN4* 5ʹ UTR and *ADH1* 3ʹ UTR at the native locus (Fig. 1a). This reporter retains Ssd1-dependence but shows higher abundance of Sun4-mNG, giving a higher signal-to-noise ratio in flow cytometry experiments. All Ssd1- mRuby2 variants tested have a similar ∼10-fold increase in red fluorescence over background (a plasmid lacking mRuby2), and slightly reduced fluorescence compared with a plasmid expressing mRuby2 alone (Fig. 4e). As expected, mNG fluorescence is lower (2-fold) in cells where wild-type Ssd1 is present, compared with empty plasmid controls (Fig. 4e), recapitulating Ssd1-dependent repression of Sun4. Ssd1 point mutants showed higher Sun4- mNG abundance compared to wild-type, with the largest increases seen for W583A and H630A, a less dramatic change for R394A and KR622-623AA and the most modest increase for K589A (Fig. 4e, Fig. S4b).

Overall, we characterized a set of mutants with strong to moderate loss-of-function phenotypes (R394A, W583A, H630A, and KR622-623AA) or with mild phenotypes (K589A) consistent across three assays: resistance to cell wall stress, inviability in a *cbk1Δ ssd1Δ* background, and repression of Sun4-mNG. These mutants show coherent phenotypes across all assays, underscoring the importance of the RNA binding function of Ssd1 for regulatory and growth phenotypes, across different genetic backgrounds and in normal and stress growth conditions.

### Recognition of at least one SBS by Ssd1 is required to repress protein abundance

We next sought to test the effects of mutating Ssd1 binding sites in the *SUN4* mRNA. Deletions or mutations to SBSs were introduced into the *SUN4-mNG* construct that has an intronless *SUN4* 5ʹ UTR, and a *ADH1* 3ʹ UTR (Fig. 1a). This construct had high expression and clear Ssd1- dependence (Fig. 1b) and has the advantage that changes to 5ʹ UTR sequences are unlikely to impact splicing. Short deletions in this construct removed the first SBS (Del-1), both SBSs (Del- 2), or both SBSs and extended neighbouring regions (Del-3) (Fig. 5a).

**Figure 5.**
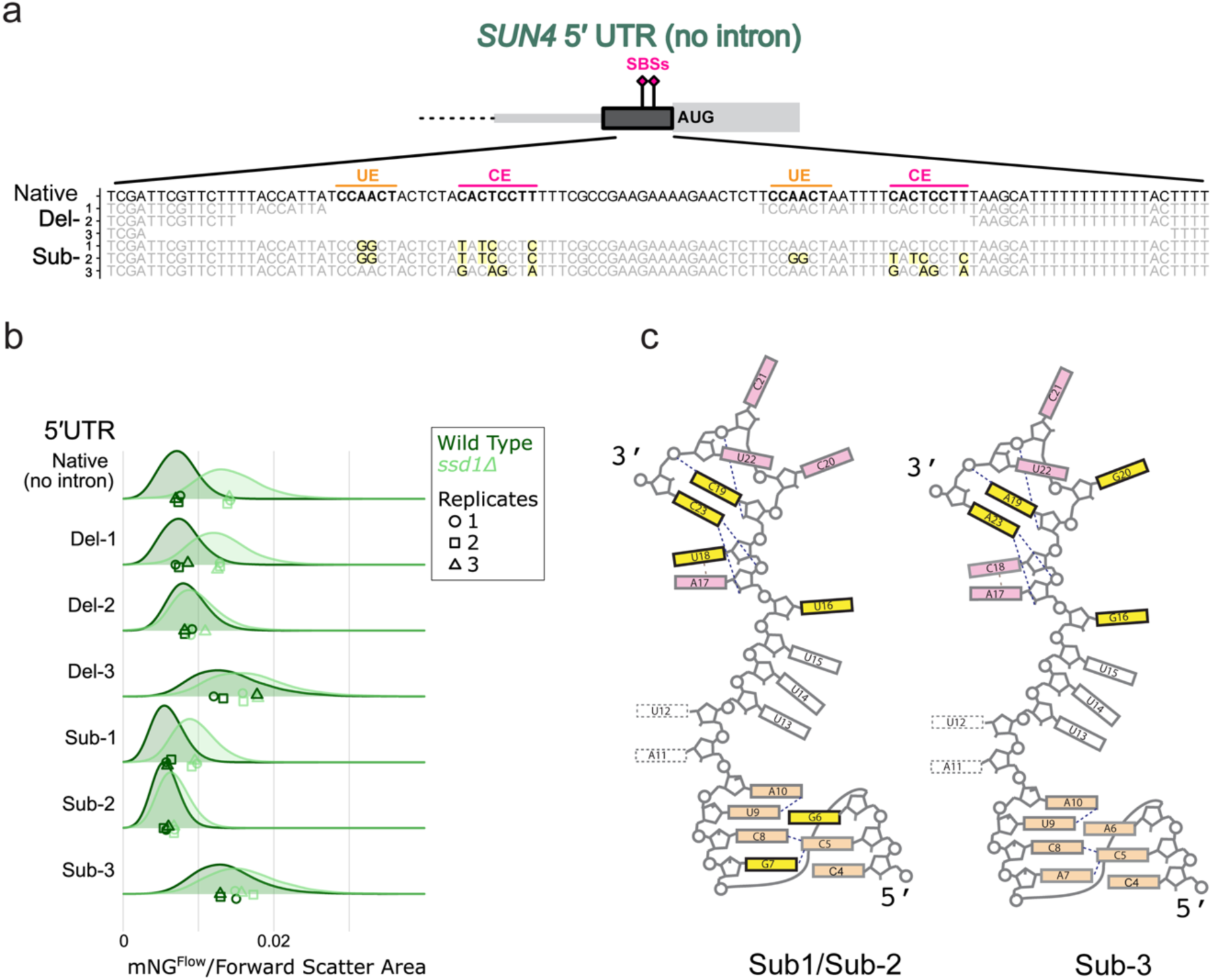
Alterations to SBSs reduce Ssd1-dependent repression of Sun4 abundance. (**a**) Overview of *SUN4* 5ʹ UTR SBS region showing tested deletions and mutations. Strains with sequence deletions are denoted ‘Del-X’ and strains with sequence substitutions are ‘Sub-X’. All mutants lack the 5ʹ UTR intron. Substitutions are highlighted in yellow. UE = SBS upstream element (CCAACU), CE = SBS CNYU element (CACTCCTT). (**b**) Fluorescence of Sun4- mNG proteins expressed from intronless native *SUN4* or Del-X/Sub-X mutants, in WT *SSD1* (dark green) and *ssd1Δ* (light green) backgrounds, measured by flow cytometry. Cell size effects are accounted for by dividing mNG fluorescence by forward scatter area. Three biological replicates, from independent single colonies, were measured. Mean values are plotted individually. Note that these data were collected in the same experiment as in Fig. 1b so the “Native (no intron)” control data are identical to that figure. (**c**) Schematic based on Fig. 3a, showing positions of RNA substitutions (yellow).

Flow cytometry showed that deleting a single SBS does not alter Ssd1-dependent repression of Sun4 protein abundance, but deleting both SBSs does (Fig. 5b). Del-1 shows a similar profile to the no-intron control, with higher fluorescence in *ssd1Δ* than in wild-type cells (Fig. 5b). In contrast, both Del-2 and Del-3 showed a loss of Ssd1-dependent repression, where wild-type and *ssd1Δ* backgrounds have overlapping fluorescence profiles. Del-3 had a higher overall abundance for Sun4-mNG in both backgrounds, showing that the extended sequences beyond the SBSs contribute to regulation of protein abundance (Fig. 5b).

To further understand how RNA sequence contributes to Ssd1-dependent regulation, we generated three sets of nucleotide substitution mutations to disrupt SBSs within the same intronless *SUN4* 5ʹ UTR parent strain (Fig. 5a). We tested mutations that only affect the first SBS (Sub-1), mutating the UE (CCAACT to CCGGCT, disrupting the hydrogen bond network in the UE and possibly destabilizing its structure) and shuffling the CE (CACTCCTT to TATCCCCT, to disrupt base-specific recognition of C16 and the bridge between U19 and U23 that latches the motif) (Fig. 5c). This was compared with equivalent mutations that affect both SBSs (Sub- 2) to test if the presence of a single native SBS retains Ssd1-dependent repression of Sun4- mNG abundance. In a third substitution, only the CE elements were disrupted (CACTCCTT to GACAGCTA, affecting base-specific recognition of C16 and C20 and the latch mechanism of U19 with U23 (Fig. 5c).

Flow cytometry showed that disruption of a single SBS maintains Ssd1-dependent repression (Sub-1), i.e. fluorescence is higher in *ssd1Δ* compared with wild-type cells (Fig. 5b). However, disrupting both SBSs (Sub-2 or Sub-3) removes the ability of Ssd1 to repress Sun4 protein abundance (Fig. 5b). The Sub-1 mutations lead to an overall reduction in protein abundance while loss of both CNYU elements in the Sub-3 mutant increases overall protein abundance (Fig. 5b). As observed in the flow cytometry experiments described above, changes to UTR sequences can have dramatic effects on protein abundance independently of Ssd1-dependent regulation (Fig 1b). As with UTR substitutions, we asked whether changes to protein abundance observed with these alterations to 5ʹ UTRs could be explained by changes in mRNA abundance using RT-qPCR. (Fig. S5a). Variation in mRNA abundance is minimal between deletion and substitution variants and between wild-type and *ssd1Δ* cells (Fig. S5a). Protein abundance and RNA abundance of *SUN4-mNG* reporters are uncorrelated (R = 0.16), indicating that changes in protein abundance are determined post-transcriptionally (Fig. S5b).

Both Del-1 and Sub-1 constructs have a single intact SBS in the 5ʹ UTR and are repressed by Ssd1 similarly to the native 5ʹ UTR, while constructs with both SBSs deleted (Del-2, Del-3) or substituted (Sub-2, Sub-3) lose Ssd1-dependent repression. We therefore conclude that one intact SBS is necessary and sufficient for Ssd1-dependent post-transcriptional repression in the context of *SUN4-mNG*.

## Discussion

What are the key determinants of RNA recognition by Ssd1 that enable regulation of protein abundance? Here, we have dissected the regulatory sequences of *SUN4*, a major Ssd1 target, using perturbations that affect only UTRs or sequences within UTRs in the native genomic context. We find that the *SUN4* 5ʹ UTR, particularly the SBSs encoded within it, drives Ssd1- dependent repression of Sun4 protein abundance. This observation differs from prior work that reported no Ssd1-dependent effects on reporter genes containing the *SUN4* 5ʹUTR (Wanless *et al*., 2014). These different observations may be attributed to the use of a plasmid-based reporter system in prior work, compared with the conservative changes we made to the native locus here. It is also possible that there are differences in the UTR definitions used previously. As the changes to Sun4 abundance that we measured were uncorrelated with mRNA levels, we can attribute Ssd1-dependent repression of Sun4 to post-transcriptional regulation.

A co-crystal structure of Ssd1 and an SBS reveals core determinants of RNA binding. The CSDs have two separate recognition surfaces, which explains why linkers of different lengths can separate upstream and CNYU elements. The CE, which overlaps with a previously described motif common to Ssd1 binding partners (Hogan *et al*., 2008), has a splayed structure where base-specific interactions drive both the unusual conformation of the RNA and readout of its sequence by Ssd1 surface residues. In contrast, the UE is primarily a structural element with only a single base directly recognised by Ssd1. This perhaps explains why UE-like motifs have not been identified outside Saccharomycetes despite CNYU motifs being identifiable outside this clade. As both elements are needed for tight binding of *Sc*Ssd1 *in vitro* it is possible that different sequences that form equivalent structures may be present in other fungal mRNAs recognised by Ssd1 homologs.

Disrupting Ssd1 recognition of RNA with structure-guided point mutations affects three well-characterized Ssd1-dependent growth phenotypes. Some point mutants i.e. R394A, W583A, H630A and KR622-623AA, fail to restore wild-type Ssd1 phenotypes of promoting cell wall stress resistance when incorporated at the native locus. Similarly, when complemented on a plasmid these mutant alleles permit growth of *cbk1Δ* cells and fail to repress Sun4 abundance. Inspection of the RNA-bound Ssd1 structure suggests why a range of phenotypes is observed. Notably, R394, R623 and H630 each contribute to base recognition and, when mutated, have stronger phenotypes in all assays. Their loss could affect both RNA binding affinity and specificity. H630, K622 and W583 each contribute to structuring the RNA in the binding site. Stronger phenotypes associated with mutations of these residues suggests that maintaining the correct conformation is also important for RNA recognition. In contrast, mutations to K585 and N370 have a lesser impact on Ssd1 function. K585 is only one of several residues that stabilise C20 binding, while N370 binds C21, which is not a highly conserved position in the RNA motif. Complementary to this, disrupting SBSs in RNA, either by deleting or altering their sequence, also leads to loss of Ssd1-dependent regulation. Together, these data indicate that the direct recognition of an SBS in the *SUN4* 5ʹ UTR by the structured C-terminal domain of Ssd1 is necessary for repression of protein abundance. RNA recognition is therefore a primary function of Ssd1.

A common pattern of Ssd1 binding to 5ʹ UTRs suggests that a similar regulatory mechanism could operate across Ssd1-bound mRNAs beyond *SUN4*. Most Ssd1-enriched mRNAs in yeast cells encode cell wall proteins and are bound via their 5ʹ UTRs, like other SUN-family genes *SIM1* and *UTH1* (Bayne *et al*., 2021). By contrast, some mRNAs are bound by Ssd1 in the 3ʹ UTR, such as the cell wall gene *HSP150/PIR2* (Bayne *et al*., 2021), and Ssd1 also regulates other groups of genes like the cyclin *CLN2* (Ohyama *et al*., 2010) that might have a different transcriptional and post-transcriptional regulatory profile.

The mRNA sequence context beyond the SBSs impacts overall expression of the encoded protein. We observed that Sun4 abundance can vary substantially with differing UTR sequences even when Ssd1-dependent repression is lost. This includes experiments where whole UTRs were replaced and where local nucleotide changes were introduced into SBSs to reduce Ssd1 binding. Indeed, two different deletions of the *SUN4* 5ʹ UTR that each remove the two encoded SBSs differ substantively in the amount of fluorescent protein produced. The results of Wanless et al. show that 3ʹ UTR sequences can also affect Ssd1-dependent regulation in different gene constructs (Wanless *et al*., 2014). Overall, this suggests that other factors such as local RNA structure or additional protein interactions may affect both total protein production from a given mRNA and its regulation by Ssd1. Indeed, Ssd1-bound mRNAs are bound by distinct complements of RNA-binding proteins (Dutcher & Gasch, 2024; Hall & Wallace, 2022; Hogan *et al*., 2008). It is therefore likely that mechanisms of post- transcriptional regulation by Ssd1 occur in cooperation with other RNA-binding proteins, across a wider set of yeast and fungal genes. This cooperation may promote diverse fungal growth programs, including in fungal pathogens.

## Materials and Methods

### Ssd1 protein expression and purification

*S. cerevisiae* Ssd1 ΔN338 used in this study was cloned, expressed, and purified as detailed in Bayne *et al*, 2022, NAR (Bayne *et al*., 2021).

### Protein-RNA complex formation

SUN4-duo-26mer RNA oligo (5’ cuuccaacuaauuuucacuccuuuaa 3’) was synthesised by Biomers GmBH and reconstituted in water to a final concentration of 1 mM. SUN4-duo-26mer RNA oligo was mixed with Ssd1 ΔN338 in a molar ratio of 1.2:1 in a total volume of 250 μl using 20 mM HEPES (pH 7.5), 150 mM NaCl and 1 mM DTT. The sample was incubated on ice for 30 min and injected into size exclusion chromatography Superdex 200 (Cytiva). The complex was eluted in buffer containing 20 mM HEPES (pH 7.5), 150 mM NaCl, 1 mM DTT. The eluted pure protein-RNA complex peak was concentrated and snap frozen.

### Crystallization and structure solution

Purified Ssd1 ΔN338 – RNA complex was concentrated to 24 mg/ml and crystallized at room temperature in sitting drops containing a mother solution of 50 mM MES (pH 5.6), 15% PEG 3350. Crystals were cryoprotected in 50 mM MES (pH 5.6), 30% PEG 3350 and cryocooled in liquid nitrogen. Data were collected at Diamond Light Source (DLS) on beamline i04 (Table 1). Data reduction was done by automated processing in DIALS version 3.8.2 (Winter *et al*, 2018) at DLS, using Xia2 version 3.8.1 (Winter, 2010). The structure was solved by molecular replacement using coordinates from PDBid 7AM1 (Bayne *et al*., 2021) with Phaser version 2.8.3 (McCoy *et al*, 2007) in CCP4i2 (Agirre *et al*, 2023; Potterton *et al*, 2018). The structure was rebuilt and refined using COOT version 0.9.8.91 EL (Emsley *et al*, 2010) and Phenix, version 1.19.2-4158 (Liebschner *et al*, 2019). Structure quality was checked using MolProbity (Williams *et al*, 2018).

### Construction of yeast strains

All strains were made in the BY4741 (S288C) background; BY4741 and *ssd1Δ* strains were sourced from the yeast gene deletion collection (Giaever *et al*, 2002) and a clean deletion of the *SSD1* ORF was verified by PCR. All oligonucleotides and gBlocks were supplied by Integrated DNA Technologies. Parts for episomal yeast expression plasmids were taken from the Yeast Modular Cloning Toolkit (MoClo-YTK; (Lee *et al*, 2015)), a gift from John Dueber (Addgene kit # 1000000061). We referred to the Saccharomyces Genome Database (Cherry *et al*, 2012) and FungiDB (Stajich *et al*, 2012) for sequence information. Cloning strategies, designed using SnapGene (GSL Biotech LLC, San Diego, CA), are shared at DOI: 10.5281/zenodo.20560340.

All yeast strains were made with lithium acetate transformation protocols (Gietz & Schiestl, 2007a, b). *SSD1* and *SUN4* point mutant and deletion yeast strains were constructed using CRISPR-Cas9 gene editing technology (Laughery *et al*, 2015; Laughery & Wyrick, 2019). Briefly, sgRNA sequences were annealed and ligated into the Cas9/sgRNA expression vector pML104, with SwaI and BclI, after growth in dam− *Escherichia coli*, and plasmid DNA was isolated using a Zymo Zyppy Plasmid Miniprep kit. For repair template design, additional synonymous mutations within the gRNA/PAM target site were included for most mutant strains, to prevent further cleavage after repair. BY4741 yeast strains were transformed and selected as described above with 1500 ng of the relevant duplexed repair template and 500 ng sgRNA expression vector. Clones were verified by PCR analysis and sequencing, then plated on 5-FOA agar to select for loss of the Cas9/sgRNA vector.

For construction of the episomal yeast expression plasmids, a low-copy centromeric (*CEN/ARS*) level 2 GFP dropout plasmid with *E. coli* and *S. cerevisiae* replication origins and a yeast *LEU2* selectable marker was first assembled via a golden gate BsaI reaction (Lee *et al*., 2015). Internal BsaI recognition sites in the GFP cassette were preserved following this reaction by omitting the final digestion and inactivation steps, facilitating replacement of the cassette in a second golden gate assembly with a fluorophore-tagged *SSD1* transcription unit (TU). For the *SSD1* TU, *SSD1* coding sequences free of BsaI sites were PCR-amplified with WT or mutagenesis primers to include MoClo YTK type 3A overhangs, and cloned into an entry vector, pYTK001, via a BsmBI golden gate reaction. The *SSD1* CDS was then cloned, along with level 1 vectors containing a type 2 *RNR2* promoter (medium expression level), type 3B *mRUBY2* fluorescent tag, and type 4 *ENO1* terminator parts, into the GFP dropout vector via BsaI golden gate assembly (Lee *et al*., 2015). Plasmid preparations were transformed into yeast strains *ssd1Δ* and yMJ011 and maintained by selection in SC-LEU dropout medium for spot assays and flow cytometry analysis.

### Yeast growth assays

We investigated the sensitivity of yeast growth to cell wall stress, by inoculating individual colonies in 3 ml YPD broth in culture tubes, and growing with vigorous shaking overnight at 30°C. To test Calcofluor White (CFW) sensitivity, overnight cultures were diluted to OD_600_ = 1.0, and 4x 10-fold serial dilutions of each strain were prepared in YPD in a UV-sterilized U- bottom 96-well plate. 5 µL of each dilution was pipetted onto YPD and YPD + 25 µM CFW plates. Plates were incubated at 30°C for 48 hours, then imaged with an Amersham™ ImageQuant 800 imaging system, on a transparent glass tray with the ‘Colorimetric OD measurement’ setting.

We investigated the effect of plasmid-expressed Ssd1 on growth of *cbk1Δ ssd1Δ* strains, by transforming plasmids into *cbk1Δ ssd1Δ* yeast and performing 10-fold serial dilutions of each transformation reaction. 5 µL of each dilution was then immediately pipetted on SC or SC-LEU plates, and plates were incubated and imaged as above.

### Yeast flow cytometry and analysis

Flow cytometry was used to quantify Sun4-mNG expression in mutant *ssd1* strains. Cells were grown to mid-log phase in 0.2-1 mL low fluorescence, synthetic complete media MES-buffered to pH 6, then acquired immediately by an LSRFortessa X-20 Cell Analyzer. For acquisition, voltages were adjusted to detect the full range of fluorescence intensities for our fluorophores, then 10000 events were acquired for each sample. 3 biological replicates, from single colonies, were acquired per strain for all experiments.

Following acquisition, FCS files were imported into R using the flowCore package (Ellis *et al*, 2026). Data were analyzed with the tidyverse package, and plots were made with ggplot2 (Wickham, 2009; Wickham & 2019). Samples were gated according to the height and width components of the forward scatter parameter (FSC-H and FSC-W, respectively), to include only singlet cells in the analysis (Fig. S1a,b). This was to ensure consistent morphology between strains, and unimodal fluorescence distributions for all samples. Fluorescence intensities from the 530/30 filter (excitation at 488 nm) and 610/20 filter (excitation at 561 nm) were used to quantify mNG and mRuby2 abundances, respectively. A strong correlation was consistently found between mNG intensity and forward scatter area (FSC-A, cell size proxy; Fig. S1b), so mNG was divided by FSC-A to account for the effect of cell size on fluorescence. Ridge / kernel density plots were generated (Wilke, 2025) to visually compare normalized fluorescence distributions between strains, and a log-linear model was fit to the FSC-A-normalized data to quantify differences between sample means. Raw data and R scripts are shared at Zenodo, DOI: 10.5281/zenodo.20560777.

### Yeast fluorescence microscopy and image analysis

Fluorescence microscopy was used to validate our flow cytometry results. Cells were grown to mid-log phase in 0.2 mL low fluorescence, synthetic complete media MES-buffered to pH 6 in a 96 well Sensoplate^™^, then diluted 4x in 1x PBS to visually distinguish cells clusters. Samples were imaged with a Nikon Eclipse Ti inverted microscope, with a 100x objective and 1.5x switching knob (150x total magnification). Images were taken in the GFPFast (LED excitation filter 470/40 nm, emission filter 520/40) of 2 fields of view per well, with the Perfect Focusing System (PFS) and 1x1 binning, and 300 ms exposures.

Raw .tiff files were imported into ImageJ (Schindelin *et al*, 2012) and split by channel, then converted to grayscale. Contrast and brightness of the brightest image were adjusted, then propagated to all other images for comparison across samples. Images were converted to 8- bit, then saved as modified .tiff files.

### Reverse transcription and quantitative PCR (RT-qPCR) analysis

RT-qPCR was performed on *SUN4-mNG* yeast strains with variant 5ʹ UTRs. RNA was extracted from 3 single-colony biological replicates per strain, for each experiment. Cells were grown to mid-log phase in 5 mL YPD aliquots, then centrifuged at 3000 g, 4°C for 2 minutes, washed in 1 mL ice-cold deionized water, and snap-frozen on dry ice. RNA was extracted from pellets following a column-based method adapted from the Zymo Quick-RNA^™^ Miniprep Kit as described previously (Haynes *et al*., 2022). Briefly, frozen pellets were resuspended in Zymo RNA Binding Buffer then lysed with a Precellys evolution touch homogenizer (Bertin Technologies). Lysates were centrifuged at high speed, then supernatants were transferred to Zymo-Spin IIICG columns and centrifuged again to remove DNA contamination. Flow-through samples were transferred to Zymo-Spin II columns and centrifuged to capture RNA. Columns were washed in Zymo DNA/RNA prep buffer then Zymo DNA/RNA wash buffer, after which RNA was eluted into RNase-free water.

RT-qPCR was performed on RNA extracts according to a protocol using Superscript IV Reverse Transcriptase, Brilliant III Ultra-Fast SYBR^®^ Green QPCR Master Mix and LightCycler 480 II instrument. Briefly, extracts were normalized by concentration to contain >1 µg RNA, then each sample treated with 2 U DNaseI to remove remaining DNA contamination. Total RNA was reverse transcribed to first strand cDNA with 100 U Superscript IV reverse transcriptase, and 7.5 µM Random Primer Mix, and diluted 20-fold prior to qPCR. A reverse transcriptase-free (−RT) control for each sample was incubated in reverse transcriptase mix as well, to control for effects of remaining DNA contamination on qPCR results. 2 U RNasin^®^ Ribonuclease Inhibitor was added to all reactions to prevent RNase contamination.

The qPCR assay was performed on 3 technical replicates of each +RT cDNA preparation, and 1 −RT replicate. Each sample was PCR-amplified with 6 previously validated primer sets, targeting the *mNG*, *SUN4*, and *SSD1* CDS, as well as the CDS of reference genes *ACT1*, *PGK1*, and *SRO9*. 2 µL diluted cDNA was mixed with 2 µL qPCR master mix (1.6 µL Brilliant III Ultra- Fast SYBR Green qPCR Master Mix, 0.4 µL primer mix at 4 µM), in individual wells of 384-well plates, with an Integra 8-channel Voyager II pipette, to ensure consistent volumes between wells. Samples were PCR-amplified on a Lightcycler 480 instrument, according to a two-step program of 40 cycles, consisting of 5 seconds at 96°C, and 10 seconds at 60°C with a green fluorescence reading.

The tidyqPCR R package was used for analysis of qPCR output (Wallace & Haynes, 2021). Briefly, +RT and −RT sample Cq values were compared to ensure that −RT were sufficiently high or absent, to indicate that DNA contamination is below or at the low end of the detection limit of the assay. Reference primer Cq values were checked for consistency across samples, then Cq values of samples amplified with *mNG*- and *SUN4*-targeting primers were normalised by calculating ΔCq against the median Cq values of the reference genes. ΔCq values were again normalised by calculating ΔΔCq values against the median of a single reference sample (WT or *SUN4* 5ʹ UTR without intron), and ΔΔCq were plotted for each biological replicate, by strain. Linear models were fit to the ΔΔCq values to quantify differences between sample means.

### Western blot

50 mL YPD cultures in 250 mL Erlenmeyer flasks were inoculated with yeast strains at OD_600_ = 0.05 and grown at 30⁰C with 190 RPM shaking, to a density of ∼ 7x10^6^ cells (OD_600_ = 0.5). To harvest cells, cultures were transferred to 50 mL falcon tubes and centrifuged for 3 minutes at 2500g. Cells were washed once in 10 mL ice-cold 1x PBS, then resuspended in 1 mL PBS and transferred to 2 mL screw-cap tubes. These were centrifuged for 1 minute at 5000g, and the supernatant was completely removed from each tube. Extracts were then prepared using an alkaline lysis method (Kushnirov, 2000); pellets were resuspended in deionized water at a concentration of ∼ 3.5x10^6^ cells/mL (normalized by OD_600_). 100 µL of each suspension was added to 100 µL 0.2 M NaOH. These were incubated at room temperature for 5 minutes, then centrifuged at >10,000g for 3.5 minutes, and supernatants were completely removed. Pellets were resuspended in 50 µL SDS-PAGE sample buffer (60 mM Tris-HCl pH 6.8, 5% glycerol, 2% SDS, 4% β-mercaptoethanol, 0.0025% bromophenol blue, 1x cOmplete^™^ Mini Protease Inhibitor Cocktail protease inhibitor) and boiled in water for 3 minutes. Lysates were centrifuged again at >10,000g for 3.5 minutes, and ∼150 µg protein from each supernatant was loaded into NuPAGE^™^ 3-8% 1 mm Tris-Acetate Mini Protein Gels, with 5 uL Color Prestained Protein Standard, Broad Range marker. Gels were run in an XCell SureLock^™^ Mini- Cell at 190 volts for ∼1.5 hours in NuPAGE^™^ Tris-Acetate SDS Running Buffer, then blotted in a Mini Trans-Blot^®^ Cell to 0.45 µm nitrocellulose membranes for 2 hours 40 minutes at 4⁰C with gentle stirring. Membranes were stained with Ponceau-S for total protein, then blocked for 1 hour in 5% milk in PBS with 0.05% Tween-20. Membranes were incubated in 5% milk/PBS-T with 1x Monoclonal ANTI-FLAG^®^ M2-Peroxidase (HRP) antibody produced in mouse for ∼18 hours at 4⁰C, then washed 3 times for 5 minutes each in PBS-T. Membranes were incubated for 2 minutes in Pierce^™^ ECL Western Blotting Substrate, then imaged with an Amersham ImageQuant^TM^ 800 using the chemiluminescence setting, with 10-minute exposure and 2x2 binning.

## Supporting information

Biomaterials

## Acknowledgements

We are grateful to staff at Diamond Light Source for assistance with data collection and to Chris Hall and SBS flow cytometry facility for help with flow cytometry data collection and advice on data analysis. We are grateful to Ivan Clark for help and advice with microscopy and to Jean Beggs for advice and support. Snigdha Manoj contributed to making plasmids.

BK and MMJ were supported by PhD studentships from the Darwin Trust of Edinburgh. WAD was supported by a PhD studentship from the Medical Research Council Precision Medicine Program (MR/N013166/1). AGC, UJ and AR were supported by a Wellcome Senior Fellowship (200898 to AGC). EWJW was supported by Wellcome via a Sir Henry Dale Fellowship (208779 to EWJW) and a Bioimaging Technology Development Award (310933). This work was carried out in the Wellcome Centre for Cell Biology that was supported by core funding from the Wellcome Trust (203149). This work utilized the Edinburgh Protein Production Facility (EPPF) and the Centre for Optical Instrumentation Laboratory funded by Wellcome Core Grants 092076 and 203149.

## Author contributions

**BK –** data production, analysis, co-wrote MS

**UJ –** data production, analysis, editing MS

**MMJ** – strain generation, assay development

**WAD -** strain generation, assay development

**AR –** data analysis

**EWJW –** conceptualisation, funding, data analysis, supervision, co-wrote MS

**AGC –** conceptualisation, funding, data analysis, supervision, co-wrote MS

## Data access

**Structure coordinates -** pdb_000032py (PDB ID 32PY)

**Raw data and R scripts -** DOI: 10.5281/zenodo.20560777

**Plasmid maps and raw images -** DOI: 10.5281/zenodo.20560340

