## Supplemental Figures for "The fungal RNA-binding protein Ssd1 represses Sun4 protein abundance through recognition of 5ʹ UTR structural and sequence elements"

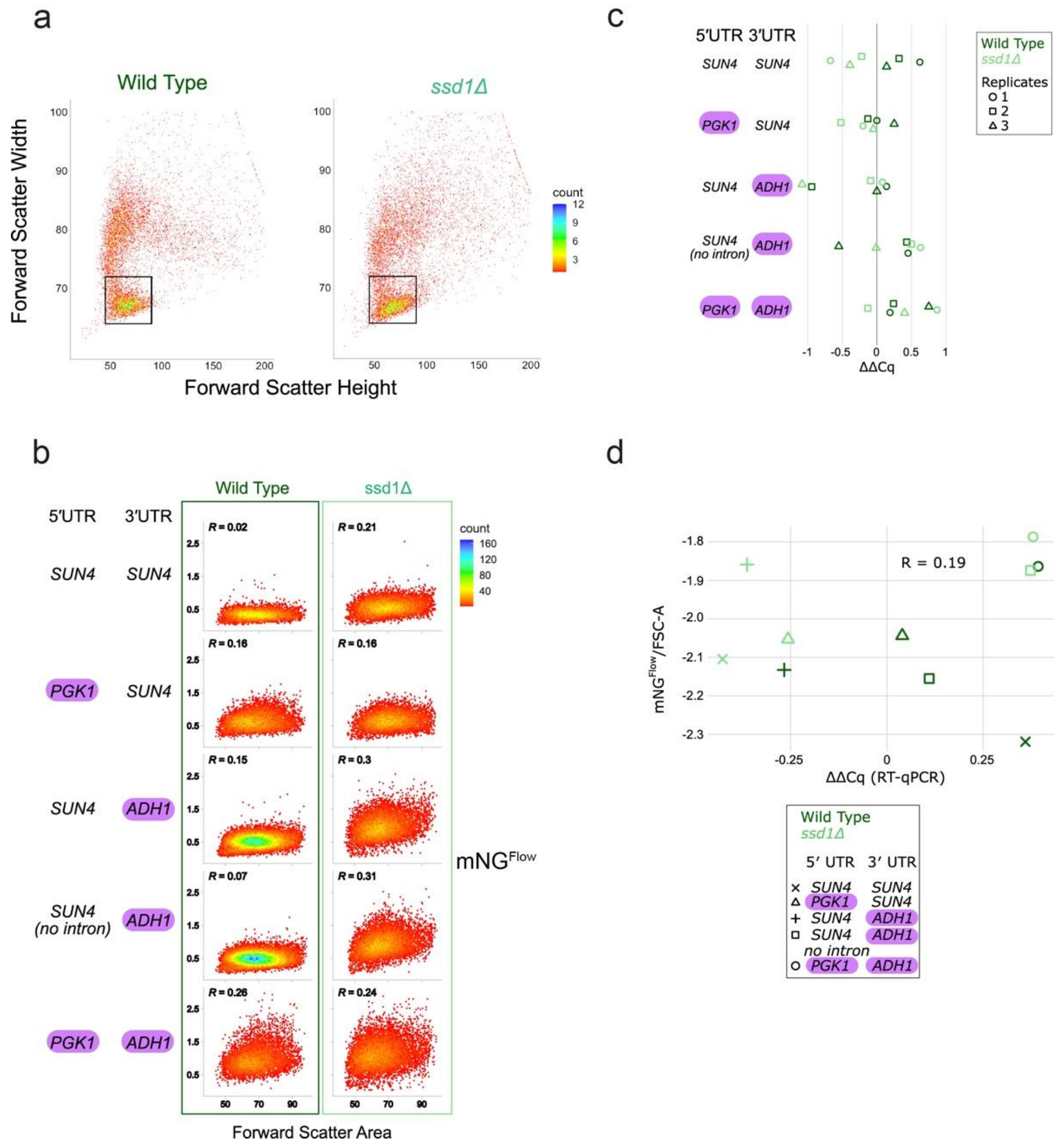


**Figure S1**. ***SUN4-mNG* mRNA abundance does not correlate withSun4 protein abundance.**

(**a**) Flow cytometry data is gated to include singlet cells. Scatterplot of forward scatter width (FSC-W) against forward scatter height (FSC-H) for WT and *ssd1Δ* strains with *SUN4-mNG* reporters. Black boxes outline cells included in downstream analysis of fluorescence intensity, for all strains. Forward scatter parameters are reported in arbitrary units x 1000. One biological replicate, from a single colony, is plotted for each strain. (**b**) Sun4-mNeonGreen signal from singlet cells shows an intermediate degree of positive correlation with forward scatter area, for *sun4* variants with replaced UTRs. Faceted scatterplot of Sun4-mNeonGreen signal from singlets against forward scatter area, for WT *SUN4-mNG* and sun4 variants with replaced UTRs, in WT and *ssd1Δ* backgrounds. Pearson's R values are reported in the upper left of each facet. Forward scatter parameters are reported in arbitrary units x 1000. Each facet includes three biological replicates, from single colonies. (**c**) ΔΔCq values are shown for native *SUN4-mNG* and variants, normalized to native *SUN4-mNG* mRNA and to 3 reference mRNAs (see methods). qPCR primers were designed to target the mNG coding sequence. Three biological replicates from independent single colonies were measured. (**d**) Scatterplot of linear model coefficients calculated from Sun4-mNG/FSC flow cytometry distributions, against coefficients calculated from RT-qPCR ΔΔCq values, for replaced-UTR variants. Shapes indicate the SUN4 construct assayed, and color indicates WT and *ssd1Δ* backgrounds. The mean values of three biological replicates, from independent single colonies, are plotted for both flow cytometry against ΔΔCq. The linear regression coefficient is stated.

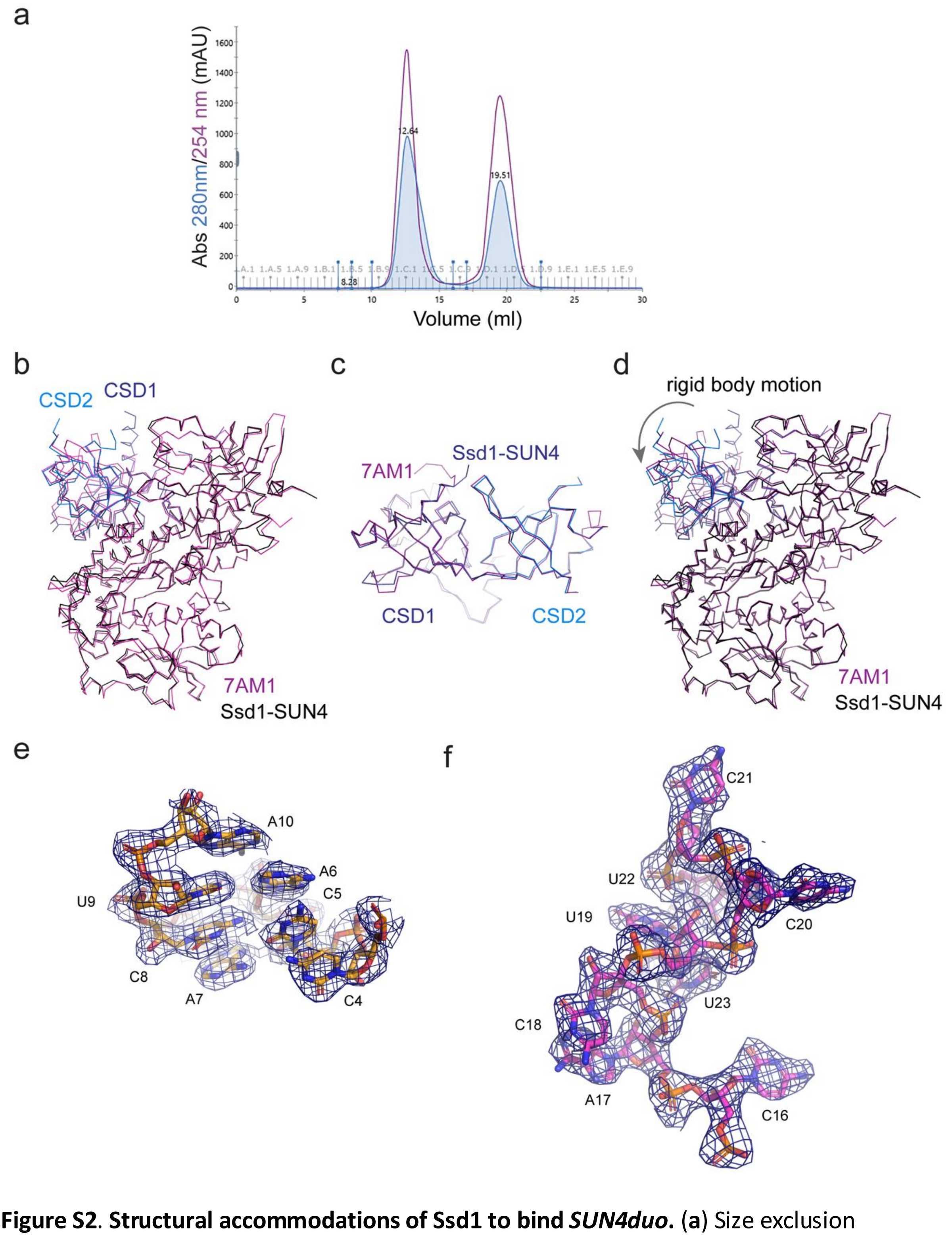


**Figure S2**. **Structural accommodations of Ssd1 to bind *SUN4duo*.** (**a**) Size exclusion chromatogram of Ssd1-*SUN4duo* complex purification. The earlier peak was pooled for crystallization. The second peak is excess RNA. (**b**) Ssd1 structure from this study(blue and black) superimposed on apo Ssd1, (7AM1, purple) using all Cα atoms. The view is the same as Fig. 2b. (**c**) CSD domains of Ssd1 (this study, dark and marine blue) superimposed on apo Ssd1. The view is similar to Fig. 2c. (**d**) As in (**b**), but with superposition over the RNB and S1 domains only. The apparent rigid body rotation of the CSDs is indicated. (**e**) Electron density of 2Fo-Fc map contoured at 1s (dark blue mesh) around SUN4 UE shown as sticks. Orientation is similar to Fig. 3b. (**f**) Electron density of 2Fo-Fc map contoured at 1$\sigma$ (dark blue mesh) around SUN4 CE shown as sticks.

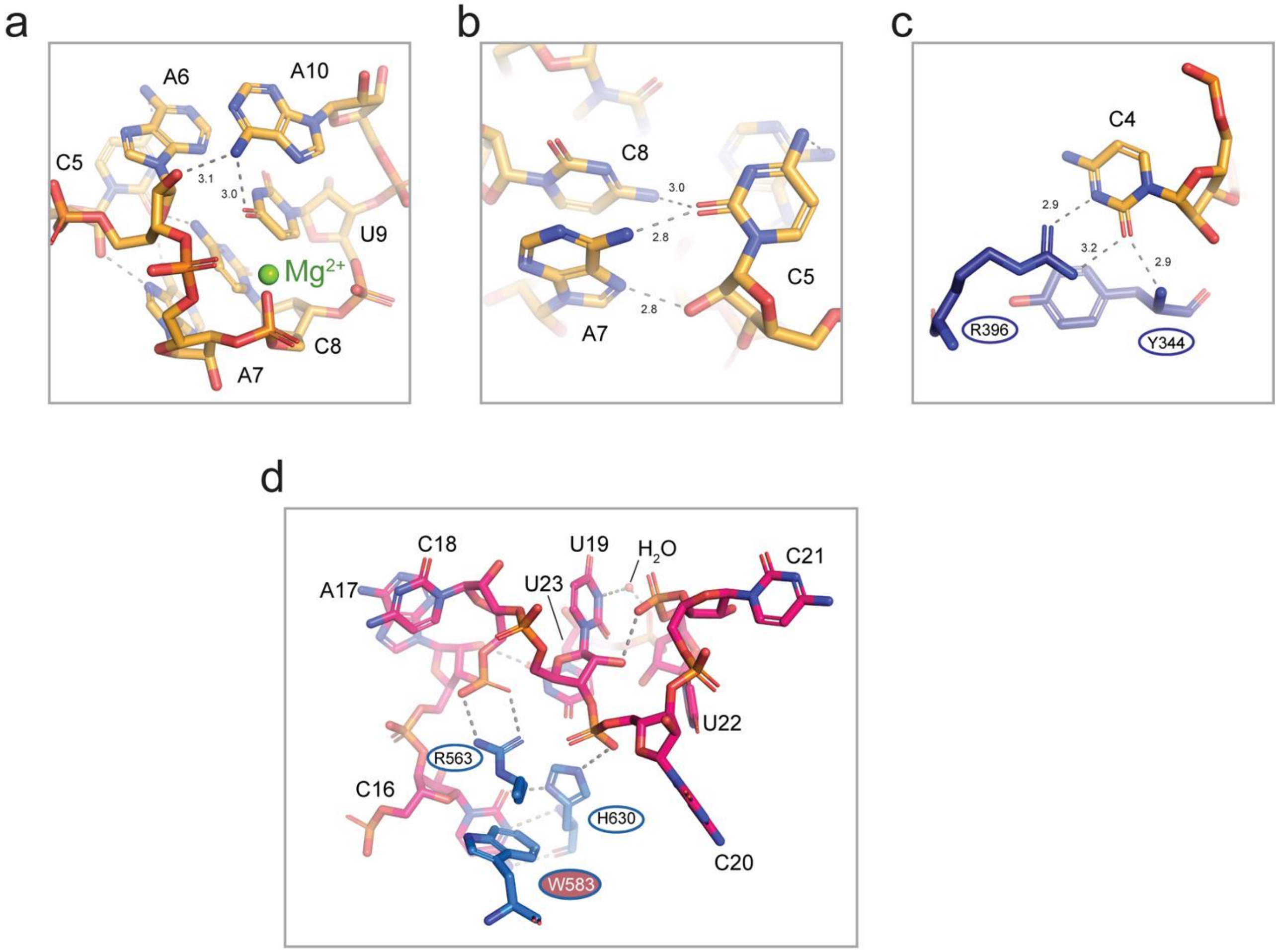


**Figure S3**. **The SBS is recognized through both structure- and sequence-specific interactions.** (**a**) Hydrogen bonding network around C5. (**b**) Hydrogen bonding network around A10. (**c**) Recognition of C4. (**d**) Backbone interactions stabilizing the flipped out A17 and C18 residues.

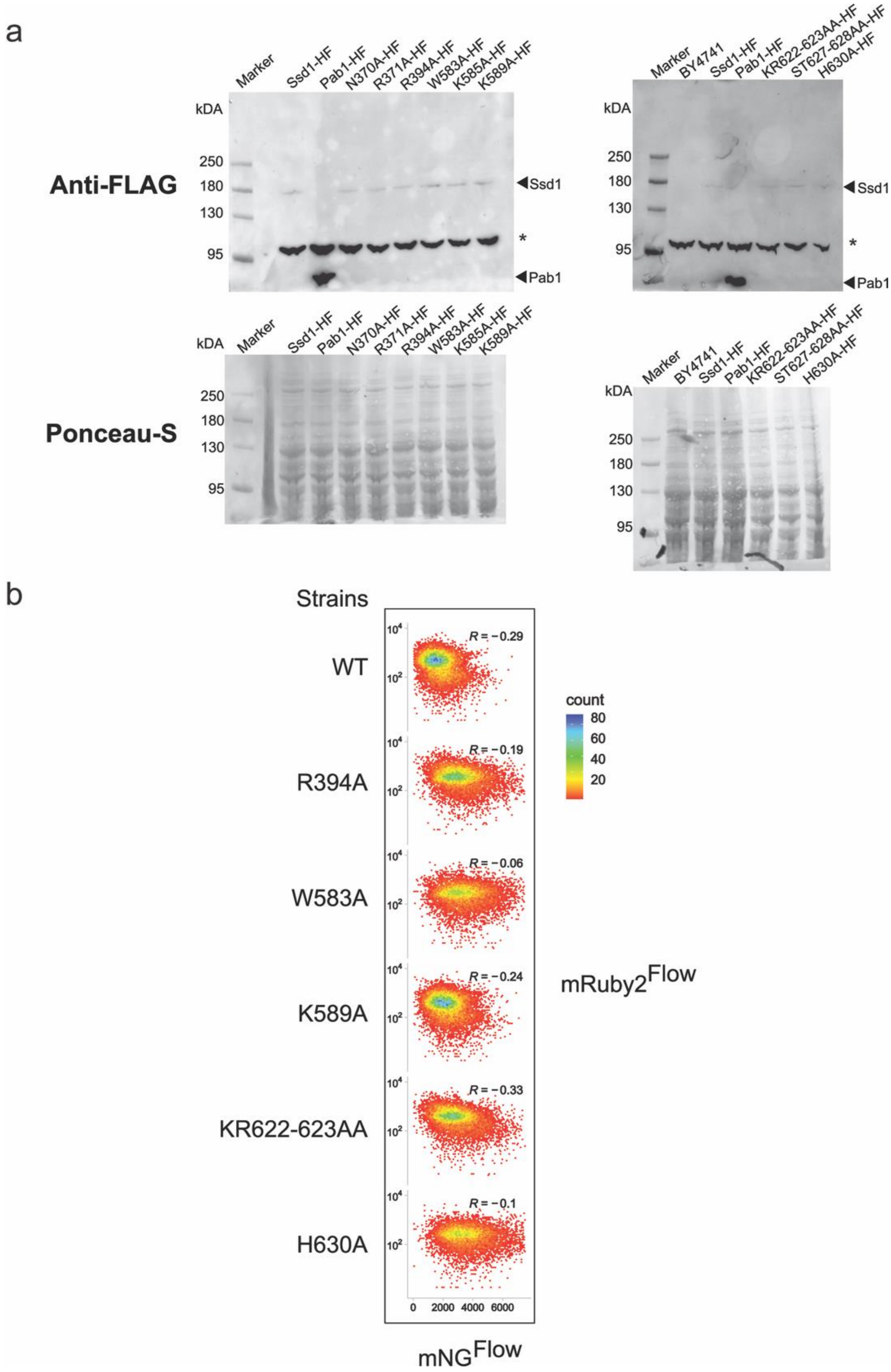


**Figure S4**. (**a**) Ssd1 RNA-binding point mutants edited at the native locus are expressed at similar levels in yeast strains. Western blots of soluble protein extracted from Ssd1 point mutant strains, in which anti-FLAG antibody was used to probe tagged versions of Ssd1-HF and Pab1-HF as a positive control. Blots were stained with Ponceau-S prior to antibody incubation, to visualise total protein. BY4741 (wild-type) was used as the untagged negative control strain. In the anti-FLAG blot, a background band at approximately 100 kb appears in all strains. (**b**) Ssd1-mRuby2 signal from singlet cells shows an intermediate degree of negative correlation with Sun4-mNG signal, for plasmid-expressed RNA-binding point mutants. Faceted scatterplot of Ssd1-mRuby2 signal from singlet cells against Sun4-mNG, for WT complementation and RNA-binding point mutants. Ssd1-mRuby2 fluorescence is displayed on a log scale, and Sun4-mNG fluorescence on a linear scale. Pearson's R values are reported in the upper right of each facet. Each facet includes three biological replicates, from independent single colonies.

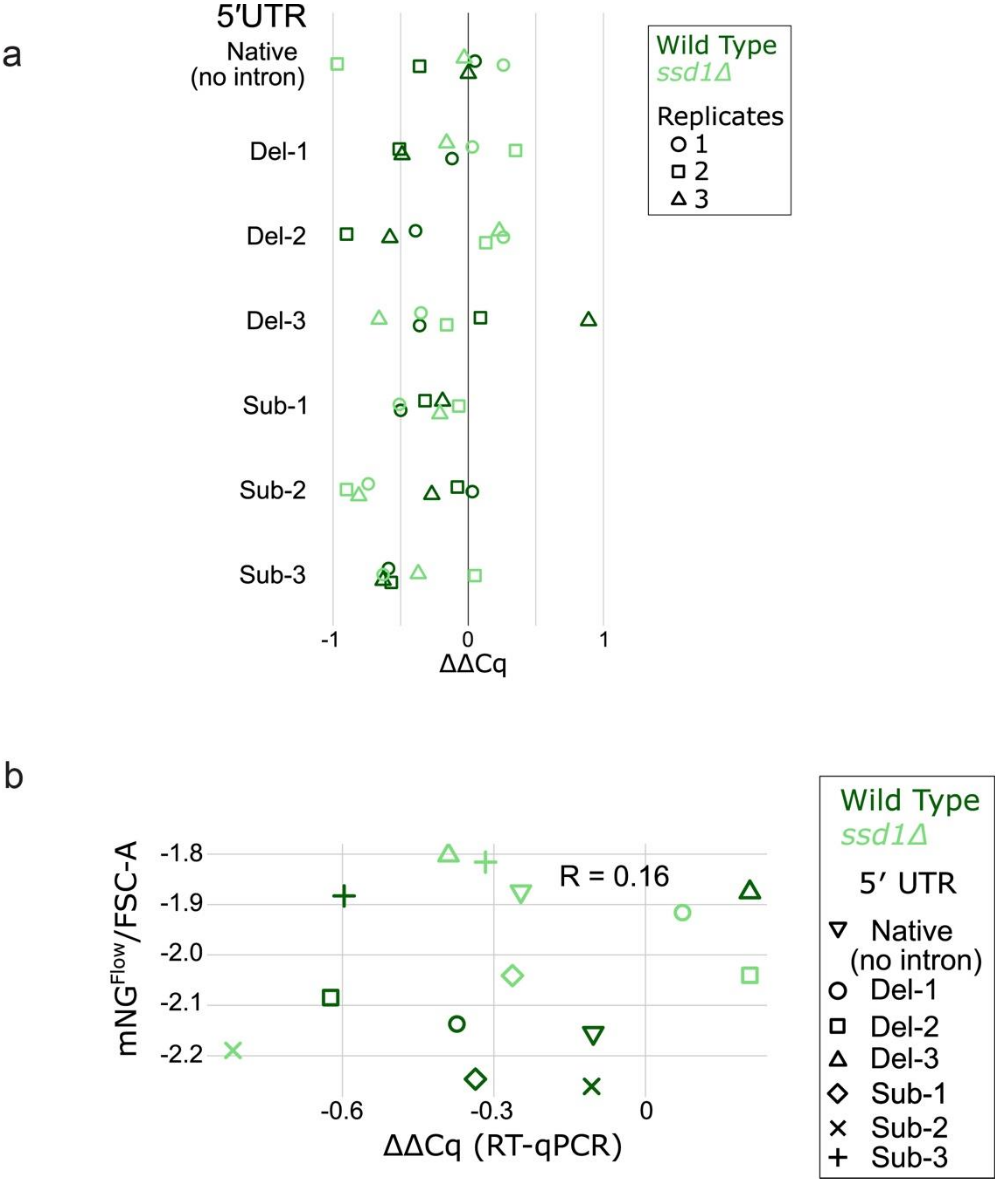


**Figure S5**. **For transcripts with native and mutated 5′ UTRs, *SUN4-mNG* mRNA abundance does correlate with Sun4 protein abundance.** (**a**) RT-qPCR ΔΔCq values for *SUN4-mNG* 5′ UTR mutants, normalized to WT (intronless) *SUN4-mNG* mRNA. qPCR primers amplify mNG coding sequence. Plotted are three biological replicates from independent single colonies. (**b**) Scatterplot of linear model coefficients calculated from Sun4-mNG/FSC flow cytometry distributions, against coefficients calculated from RT-qPCR ΔΔCq values, for WT intronless Sun4-mNeonGreen and Del-X/Sub-X mutants. Shapes indicate the *SUN4* construct assayed, and colour indicates WT (dark green) and *ssd1Δ* (light green) backgrounds. The means of three biological replicates, from independent single colonies, are plotted for both flow cytometry and ΔΔCq. The linear regression coefficient is stated.
